# Identification and Characterization of Human Activation-Induced ChAT^+^CD4^+^ T Cells

**DOI:** 10.1101/2021.04.27.441632

**Authors:** L Tarnawski, AL Gallina, EJ Kort, VS Shavva, Z Zhuge, D Martínez-Enguita, M Weiland, AS Caravaca, S Schmidt, FH Wang, A Färnert, E Weitzberg, M Gustafsson, M Eberhardson, H Hult, J Kehr, SG Malin, M Carlström, S Jovinge, PS Olofsson

**Affiliations:** Laboratory of Immunobiology, Center for Bioelectronic Medicine, Division of Cardiovascular Medicine, Department of Medicine, Solna, Karolinska Institutet, Karolinska University Hospital, Stockholm, Sweden; DeVos Cardiovascular Program, Van Andel Research Institute and Fredrik Meijer Heart and Vascular Institute/Spectrum Health, Grand Rapids, MI; Department of Physiology and Pharmacology, Karolinska Institutet, Stockholm, Sweden; Department of Physics, Chemistry and Biology, Linköping University; Pronexus Analytical AB, Bromma, Sweden; Department of Medicine, Solna, Karolinska Institutet, Stockholm, Sweden; Department of Mathematics, Royal Institute of Technology, Stockholm, Sweden; Cardiovascular Institute, Stanford University, Palo Alto, CA, U.S.A.; Institute of Bioelectronic Medicine, Feinstein Institutes for Medical Research, Manhasset, New York, U.S.A.

## Abstract

Vasodilation is a cornerstone of inflammation physiology. By regulating vasodilation and tissue entry of T cells, CD4^+^ T lymphocytes expressing choline acetyltransferase (ChAT), a key enzyme for biosynthesis of the vasorelaxant acetylcholine (ACh), critically link immunity with vascular biology in mice. However, the characterization of primary human ChAT^+^ T cells remained elusive. Here, we identified human ChAT^+^ T cells and report that *ChAT* mRNA was induced by activation. Functional studies demonstrated that T cell-derived ACh increased muscarinic ACh-receptor dependent NO-synthase activity and vasorelaxation. Further, single-cell RNA-sequencing revealed *ChAT*^+^*CD4*^+^ T cells in blood from patients with severe circulatory failure and a high relative frequency of *ChAT*^+^*CD4*^+^ T cells correlated with better 30-day survival in this cohort. Our findings provide the first insights into ChAT biology in primary human T cells, linking ChAT^+^ T cells with vasorelaxation as well as survival in a cohort of critically ill patients.

## Introduction

Vasodilation is a key physiological mechanism of inflammation involved in the homeostasis of immune responses ^1,2^. It plays a vital role in tissue entry of immune cells, and impaired vasodilation with reduced extravasation of appropriate immune cells weakens the anti-viral defense in experimental animals ^3^. A well-established mechanism for regulation of vasodilatation and blood flow is muscarinic acetylcholine (ACh) receptor mediated nitric oxide (NO)-release from vascular endothelial cells. This mechanism promotes relaxation of vascular smooth muscle cells, improves local blood perfusion and facilitates immune cell access to tissues ^4–6^. In clinical studies, low plasma ACh-levels and reduced endothelial release of the vasorelaxant NO were associated with increased mortality in the critically ill ^7,8^. However, the cellular source for ACh in blood has long been uncertain, particularly as most arteries and the endothelium lack direct cholinergic innervation ^9^. Recent discoveries in experimental models show that ACh-producing choline acetyltransferase (ChAT)^+^ T cells promote tissue entry of antigen-specific T cells as well as virus eradication as a result of vasodilation ^3^. ChAT^+^ CD4^+^ T cells also regulate blood pressure in mice, thus linking T cells, anti-microbial immunity, and vascular biology ^3,10–12^.

Several reports discuss the so called non-neuronal cholinergic system in human immune cells, such as mononuclear leukocytes ^13–16^ and immortalized human T lymphocyte cell lines ^17^, but their functional relevance is unclear. Considering the vital functions of ACh and ACh-releasing T cells in experimental animals, it becomes important to understand whether primary human T cells have the capacity to biosynthesize ACh ^13,14,18^ and promote vasodilatation.

Accordingly, we proceeded to identify and characterize primary human ChAT^+^CD4^+^ T cells. Our findings show that ChAT is induced by activation and regulated by GATA3 in a subset of blood-derived human T cells. Human *ChAT*^*+*^ T cells released ACh that promotes vasodilation, and their relative frequency was linked with survival in a cohort of critically ill patients. These results represent the first insights into ChAT biology in primary human T cells and open new avenues for research into this link between T cell immunity and vascular biology.

## Results and Discussion

### T cell activation increased *ChAT* levels and promoted ACh release from primary human lymphocytes

To investigate human T cell *ChAT* mRNA expression we turned to blood-derived human T cells from healthy individuals. Activation of human T cells using anti-CD3/CD28 antibodies significantly increased *ChAT* mRNA levels (Figure 1A). Flow cytometry analysis of cultured T cells after 72 h of activation showed that a majority of the large (defined as SSC/FSC high) cells stained positive for the activation marker CD25 (Figure 1B, S3). *ChAT* mRNA was only detected in the population of large predominantly activated T cells (Figure 1C). These findings are in line with results from experimental animal studies and human cell lines where T cell *ChAT* mRNA levels were augmented by T cell activation ^11,17,19^ and significant induction of T cell ChAT was observed only after several days of infection in mice ^3^.

**Figure 1:**
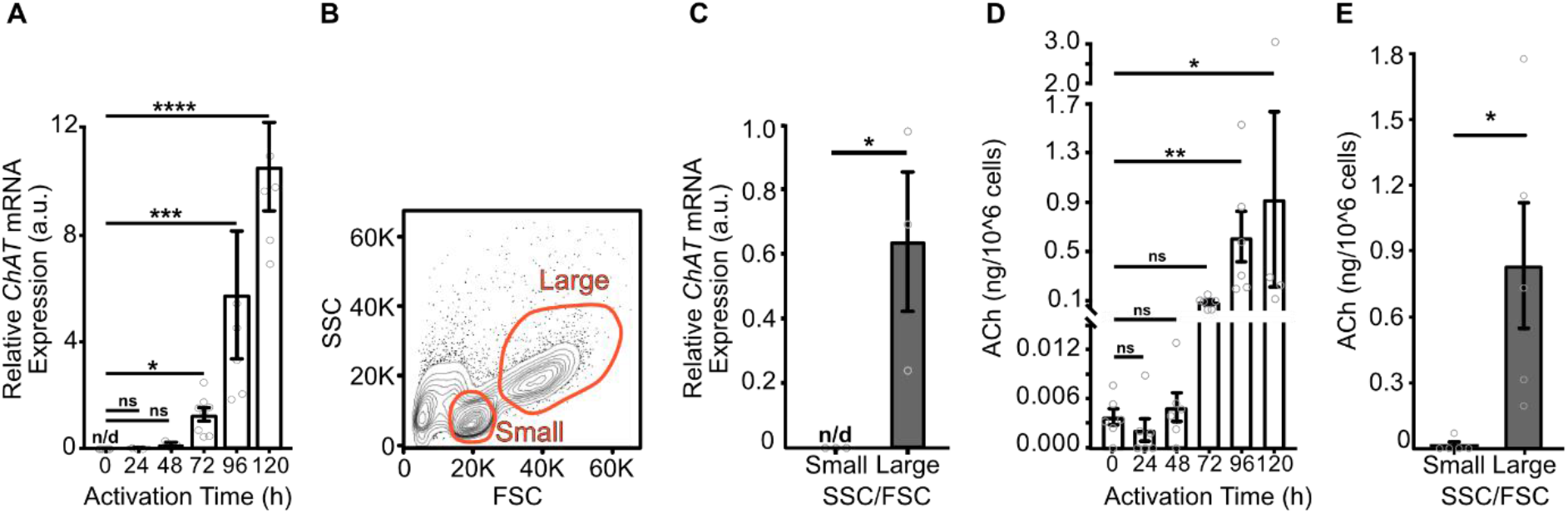
T cell activation promotes *ChAT*-mRNA and ACh release from primary human lymphocytes. **(A)** *ChAT*-mRNA was quantified by qPCR in activated primary human (n=3-8) lymphocytes harvested at indicated time points. Bars show mean ± SEM, normalized to *PPIA* and *B2M*, and expressed relative to the *ChAT*-mRNA level at 72 h (Kruskal–Wallis test, post hoc Dunn’s multiple comparisons test). **(B)** Gates used for the isolation of activated primary human lymphocytes (96 h). Size selection: Small (SSC/FSC-Low) and Large (SSC/FSC-High). **(C)** *ChAT* mRNA was measured by qPCR in FACS isolated small and large activated (96 h) primary human (n=3) lymphocytes. Bars show mean ± SEM, normalized to *PPIA*, and graphed relative to *ChAT* in large cells (Two-tailed, unpaired Student’
ss *t* test). **(D)** ACh released following activation of human (n=6) lymphocytes in vitro. The supernatants were analyzed for ACh concentration using mass spectrometry. Bars represent mean ng ACh level ± SEM per 10^6^ T cells (Kruskal–Wallis test, post hoc Dunn’s multiple comparisons test). **(E)** ACh released from large and small human (n=3) activated lymphocytes isolated using FACS. The supernatants were analyzed for ACh using mass spectrometry. Bars represent mean ACh concentration ± SEM per 10^6^ T cells (Two-tailed, unpaired Student’s *t* test). n/d – not detected, ns – not significant, *p<0.05, **p<0.01, ***p<0.001.

As ChAT is a key enzyme for ACh biosynthesis ^20^, we investigated whether human *ChAT*^+^ T cells release ACh. Activation of CD3^+^ cells using anti-CD3/CD28 antibodies significantly increased the amount of ACh in the supernatant (Figure 1D). Importantly, there was a strong correlation between *ChAT* mRNA levels and ACh release (Figure S1), and ACh was primarily released from the *ChAT*-expressing population of large T cells (Figure 1E).

The observations that activated human T cells express ChAT and release ACh suggest that ACh biosynthesis by human ChAT^+^ T cells may play a role in human inflammation and blood vessel physiology similar to what has been observed in animals ^3,11,19^. This is of particular interest because vasodilation by mouse ChAT^+^ T cells is required to control chronic viral infection ^3^. Accordingly, we next investigated whether T cell-conditioned supernatants promoted vascular relaxation.

### T cell-derived ACh induced endothelial nitric oxide synthase activity and promoted vasorelaxation

ACh promotes vasorelaxation by endothelial muscarinic ACh-receptor-dependent NO-release through activation of endothelial nitric oxide synthase (NOS) ^4–6^. In experimental animal models, ACh-producing *ChAT*^+^ T cells regulate blood pressure and blood vessel diameters in inflammation ^3,10^. Using an aortic vascular tension assay ^21^ we measured whether T cell-conditioned supernatants promoted vascular relaxation. Exposure of explanted and pre-constricted aortic rings from mice to these T cell-conditioned supernatants resulted in significant vasorelaxation, as compared to vehicle control (Figure 2A, Figure S2). The effect was abolished by pre-treatment with atropine, a muscarinic receptor blocker (Figure 2A). This observation shows that T cell-derived ACh acts through muscarinic ACh receptor in sufficient amounts to promote arterial relaxation. In line with these observations, exposure of human umbilical vein endothelial cells (HUVECs) to T cell-conditioned supernatants resulted in increased accumulation of NO related species in the culture over time, as compared to vehicle exposure (Figure 2B). Increased NO-release is mediated by activation of NOS in vascular endothelial cells. Exposure of human endothelial EA.hy926 cells to T cell-conditioned supernatant increased their NOS activity, as measured by the production of NO related species (Figure 2C) and the effect was abolished by atropine, a muscarinic receptor blocker (Figure 2C).

**Figure 2:**
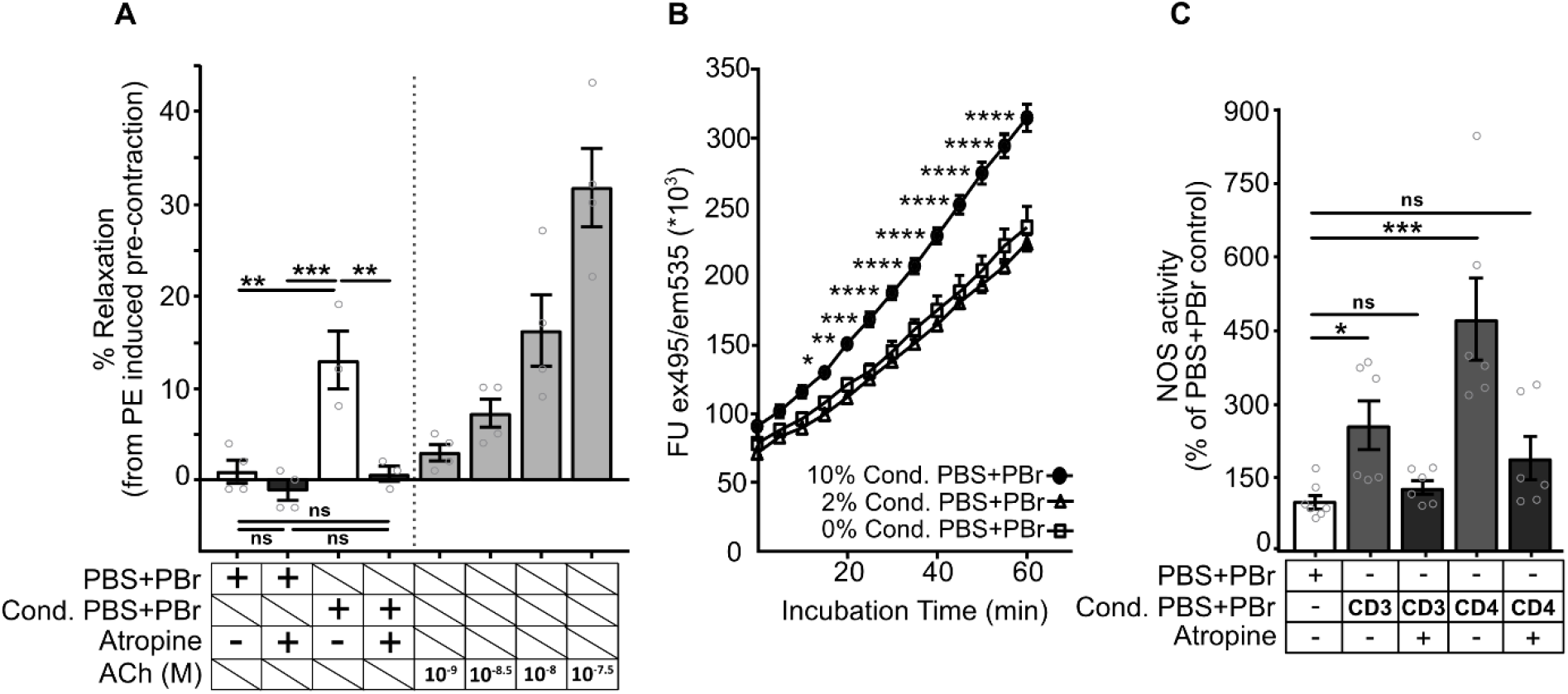
ACh release in human lymphocytes induced by activation promotes arterial relaxation. **(A)** Vascular relaxation of vessels after exposure to activated human lymphocyte-conditioned PBS. Bars show percentage of relaxation compared to maximal PE-induced pre-contraction (mean ± SEM). (left) Pre-contracted mouse aortas were exposed to vehicle (n=4) or conditioned PBS (n=3), with or without atropine pre-treatment (One-way ANOVA, Tukey’s multiple comparisons test). (right) Pre-contracted mouse aortas were exposed to ACh at indicated concentrations. **(B)** Release of NO related species from human umbilical vein endothelial cells following exposure to vehicle (n=3) or activated human lymphocyte-conditioned supernatants (n=6) at 10% or 2%. NO related species were measured by 4-amino-5-methylamino-2’,7’-difluorofluorescein fluorescence intensity assay. The fluorescence intensity is plotted as mean ± SEM (Two-way ANOVA, Dunnet’s post hoc test). **(C)** NOS activity in EA.hy926 endothelial cells following exposure to activated human CD3^+^ (n=2) or CD4^+^ (n=3) lymphocyte-conditioned PBS, with and without atropine. The cellular NOS activity was measured using colorimetric Griess assay. Bars represent NOS activity as percentage of vehicle control ± SEM (Kruskal–Wallis, Dunn’s multiple comparisons post-hoc test). ns – not significant. *p<0.05, **p<0.01, ***p<0.001.

Our observations show that human T cell-release of ACh to the extracellular space is sufficient to promote arterial relaxation through muscarinic ACh receptor activation. The cell density and resulting ACh concentration in these experiments are in a physiological range relevant to human blood ^8,22,23^, suggesting that ACh released by activated human T cells could be a source for blood ACh and sufficient to promote local vascular relaxation in vivo. Should further studies corroborate that human ChAT^+^CD4^+^ T cells indeed promote vasodilation, it would be conceivable to use these insights for therapeutic purposes in patients that require local vasodilation to improve regional circulation and tissue entry of immune cells. However, to initiate these new avenues of clinical research, it is imperative to characterize human ChAT^+^ T cells and understand the mechanisms behind *ChAT* mRNA induction. Thus, we proceeded to investigate *ChAT-*expression across different human T cell populations.

### *ChAT* is regulated by the Th2-associated master regulator GATA3

ChAT^+^ T cells in mice were initially described as CD3^+^CD4^+^CD44^hi^CD62L^low^ of an effector/memory phenotype ^11^. Subsequent studies have shown that multiple lineages of murine T cells have the capacity to express ChAT, including in effector, memory, follicular helper, and intestinal Th17-like T cells ^3,24^. Using CD4^+^ positive MACS selection of human blood-derived T cells, we observed that ChAT mRNA expression was significantly higher in CD4^+^ cell enriched as compared to CD4^+^ depleted T cells after 96 h of activation (Figure 3A). Congruently, ACh release was higher from FACS isolated CD4^+^, compared to CD8^+^ T cells (Figure 3B). Thus, we focused on the CD4^+^ T cell population.

**Figure 3:**
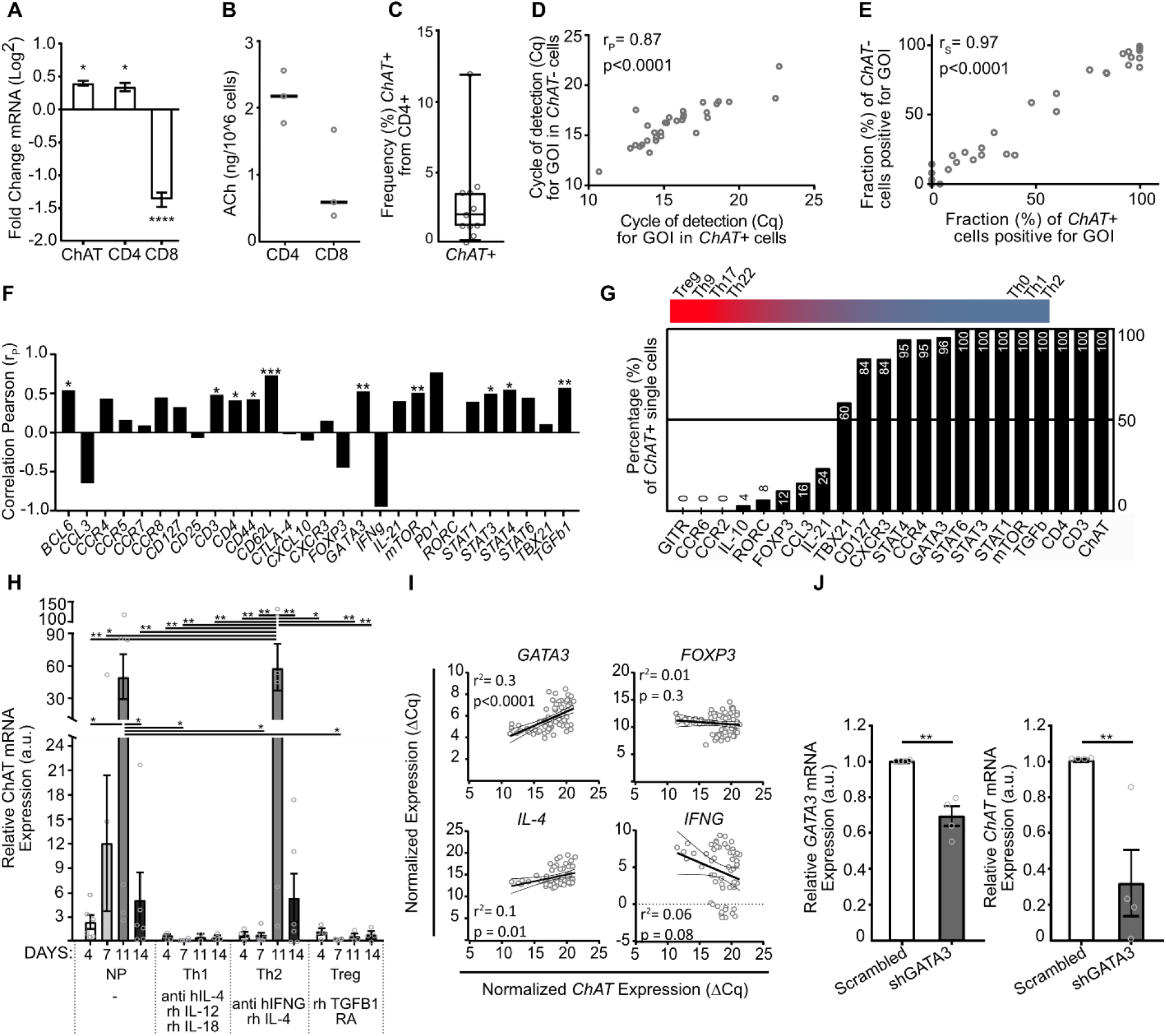
*ChAT* mRNA expression is regulated by the Th2-associated master regulator GATA3. **(A)** mRNA levels in activated CD4^+^ lymphocytes. Bars show mean ± SEM, normalized to *PPIA*/*B2M* and expressed as fold change compared to CD4 depleted cells (One-way ANOVA, Dunnett’s multiple comparisons test). **(B)** ACh released from activated CD4^+^ and CD8^+^ cells. The supernatants were analyzed for ACh using mass spectrometry. Lines: median ng per 10^6^ cells. **(C)** Frequency of *ChAT*^*+*^ cells within CD4^+^ cells (n=1631) following FACS isolation for Fluidigm Biomark single cell gene expression profiling. *ChAT* expression was measured by qPCR. The boxes indicate the interquartile range, the central bar indicates the median, and the whiskers indicate the 5^th^ to 95^th^ percentile range. Circles show *ChAT* frequency in individual donors. **(D-E)** Correlation between *ChAT*^*+*^*CD4*^*+*^ (n=25) and *ChAT*^*-*^*CD4*^*+*^ (n=153) lymphocytes: **(D)** mRNA levels and (E) fraction of cells expressing gene of interest (GOI). Each dot represents one GOI. Pearson r (r_P_), Spearman r (r_S_). **(F)** Correlation of *ChAT* and GOI-expression within activated single *ChAT*^+^*CD4*^*+*^ lymphocytes. Bars represent Pearson r (r_P_) correlation coefficients. **(G)** Frequency of GOI expression within activated single *ChAT*^+^*CD4*^+^ cells. The bars show the percentage of *ChAT*^+^*CD4*^*+*^*CD8*^-^ cells (n=19-25) that express GOI. **(H)** Relative *ChAT* mRNA levels during CD4^+^ lymphocyte polarization. mRNA was isolated at 4, 7, 11, and 14 days and mRNA levels were measured in non-polarized (NP) (n=6), T-helper 1 (n=2-6), T-helper 2 (4-6), and T-regulatory (n=4-6) polarizing conditions. Bars show mean ± SEM, normalized to *PPIA*/*B2M*, and plotted relative to NP at 4 days (Two-way ANOVA, Turkey’s post hoc test). **(I)** Correlation of *ChAT* to *GATA3, IL-4, IFNG*, and *FOXP3* during CD4^+^ lymphocyte polarization. Circles represent threshold values normalized to *PPIA*/*B2M*. Dotted line: 95% confidence interval (Linear regression). **(J)** mRNA levels of *GATA3* and *ChAT* after shRNA-mediated *GATA3* knockdown. Activated primary human (n=4) CD4^+^ lymphocytes were transfected with scrambled or shRNA targeting GATA3 mRNA. Bars show mean ± SEM, normalized to *PPIA*/*SDHA/YWHAZ*, relative to scrambled shRNA (Two-tailed, unpaired Student’
ss *t* test). *p<0.05, **p<0.01. ***p<0.001. ****p<0.0001

Next, we studied the phenotype of ChAT^+^CD4^+^ T cells. Anti-ChAT antibodies validated for staining of cholinergic neurons did not show sufficient specificity in activated primary human T cells (data not shown), which precluded use of standard flow cytometric analysis. Accordingly, we proceeded to study *ChAT*+ T cells using single cell mRNA analysis. In single activated *CD4*^+^ T cells isolated by FACS (Figure S3), an average of 2% (range 0 - 12%) of *CD4*^+^ cells within each donor expressed *ChAT* as measured by qPCR (Figure 3C). Biomark HD Fluidigm was used for single-cell analysis of 38 genes commonly associated with defined immune cell phenotypes ^25^ (Table S1) in *CD3*^*+*^*CD4*^*+*^ cells with (n=25) and without (n=153) detectable *ChAT* mRNA. There was a significant correlation of detection thresholds for the selected transcripts between *ChAT*^+^ and *ChAT*^-^ *CD4*^+^ T cells (Pearson r (r_P_) = 0.87), and none of the transcripts were significantly enriched in either of the populations (Spearman r (r_S_) = 0.97) (Figure 3D, E). Hence, these 38 transcripts failed to segregate in vitro*-*induced *ChAT*^+^ from *ChAT*^-^*CD4*^+^ T cells, supporting that ChAT expression can be induced in multiple activated T cell subtypes ^3,24^. There was a significant positive correlation between *ChAT* mRNA levels and the key regulators of T cell differentiation *BCL6, CD44, CD62L, MTOR, GATA3, STAT3, STAT4* and *TGFB1* within the in vitro-induced human *CD4*^+^ *ChAT*^+^ T cells (Figure 3F, Figure S4A). Additionally, virtually all *ChAT*^+^*CD4*^+^ T cells expressed transcripts associated with T helper (Th) 0, Th1, and Th2 subsets, but the frequency of transcripts commonly associated with Treg, Th9, Th17 and Th22 cell subsets was low (Figure 3G). To further investigate the regulation of ChAT gene expression, we used *in silico* PWMScan ^26^ to search for potential transcription binding motifs within the *ChAT* promoter. For the key regulators of T cell differentiation identified by the single cell Fluidigm analysis (Figure S4A, Figure 3F, G), potential binding sites were predicted for *STAT1, STAT4, TBX21, FOXP3*, and *GATA3* but not for *STAT3, STAT6, or BCL6* (Figure S4B).

In light of these observations, we next investigated the induction of *ChAT* in the context of T cell polarization. In mice, in vitro polarization of T cells augmented ChAT expression in Th0, Th1, and Th2 cells, but not in Th17 and Treg cells ^19^. Polarization of human T cells towards Th1 or Treg phenotypes did not significantly induce *ChAT* expression beyond four days of culture (Figure 3H). In contrast, *ChAT* mRNA was significantly induced in non-polarizing and Th2 polarizing conditions (Figure 3H). Moreover, in the non-polarizing condition, *ChAT* mRNA significantly correlated with the Th2-associated master regulator ^27,28^ *GATA3* (p<0.0001, r^2^=0.3), but not with *IL-4, FOXP3*, or *IFNG* (Figure 3I). Based on these observations and to study mechanism, we performed stable lentiviral transfection of human CD4^+^ T cells with shRNA targeting GATA3. ShRNA-mediated *GATA3* knock-down significantly suppressed both *GATA3* and *ChAT* mRNA as compared with scrambled shRNA controls (Figure 3J). Together these results show that *ChAT* in human CD4^+^ T cells, while not restricted to a particular subset, is induced by a mechanism that involves T cell activation and augmented by GATA3.

### Primary human CD4^+^ T cells in a cohort of critically ill patients express *ChAT*

We next investigated ChAT^+^CD4^+^ T cells in human clinical samples. *ChAT* mRNA was not detected in single *CD4*^+^ T cells in publicly available single cell immune profiling datasets of healthy control individuals (n=11) (Figure 4A), consistent with our observations in blood-derived human T cells from healthy individuals (Figure 1A, 0h). Since ChAT was induced by prolonged T cell activation (Figure 1A), we proceeded to investigate *ChAT* in critically ill patients with circulatory failure in imminent need of Veno-Arterial Extra Corporeal Life Support (VA-ECLS) (n=33). Patient demographics and causes of admission are shown in Table 1 and Table S2. Levels of pro-inflammatory cytokines in plasma were elevated in relation to published reference values ^29^ as shown in Table S3, indicating ongoing inflammation. All investigated critically ill patients in this cohort (n=33) had detectable *ChAT* ^+^*CD4*^*+*^ T cells in blood by single cell RNA sequencing (Figure 4A). The median relative frequency of *ChAT*^+^ cells in *CD4*^+^ T cells in the patients was 46.2 % (range 3.5 - 91 %) (Figure 4B).

**Figure 4:**
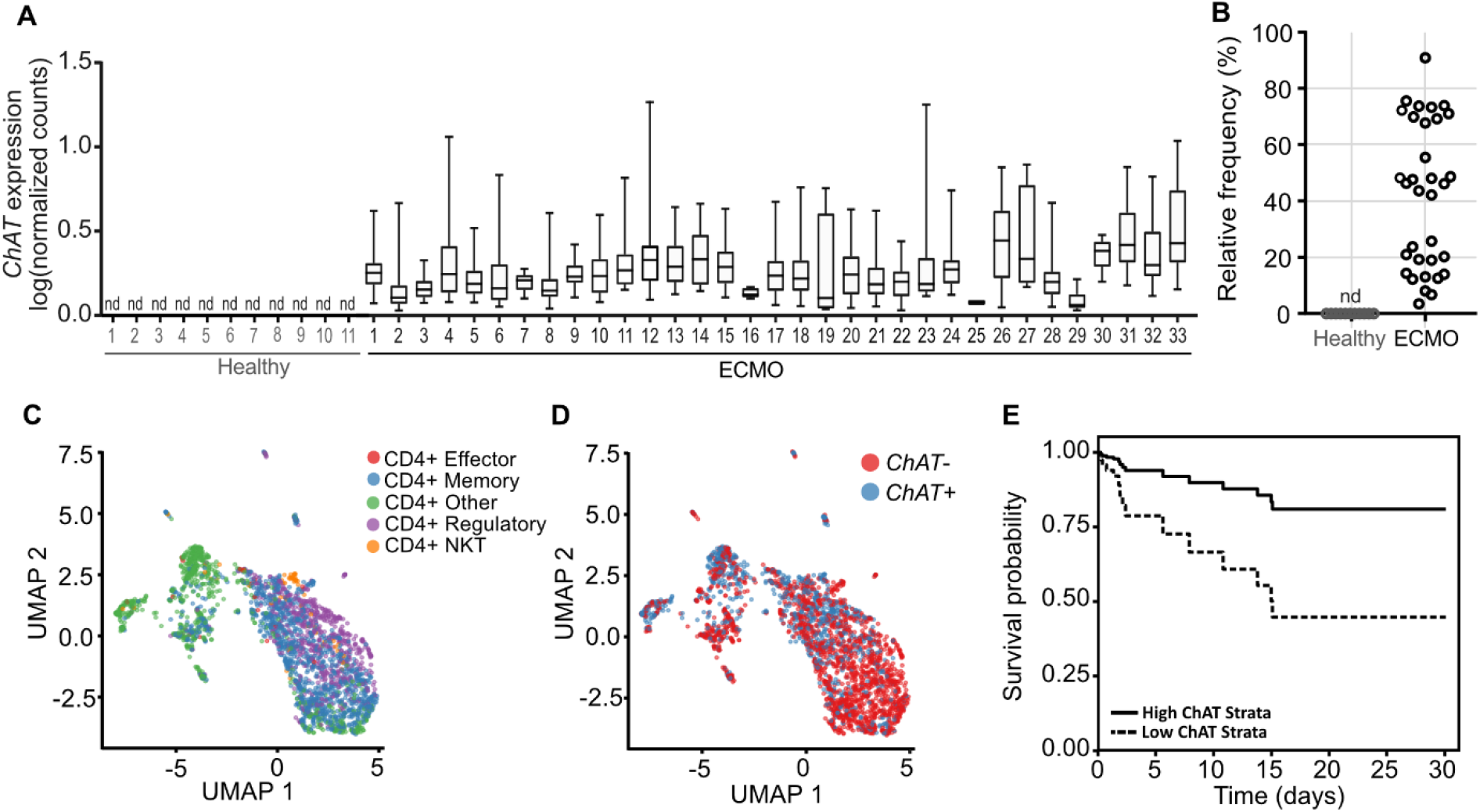
Primary human CD4^+^ T cells express *ChAT*. **(A)** Box plots indicating *ChAT* expression in single cell transcriptomic analysis of peripheral blood CD4^+^ cells from patients in circulatory failure “ECMO” (n=33) and publicly available single cell immune profiling datasets of healthy control individuals (n=11). Expression levels are represented as log normalized counts. The boxes indicate the interquartile range, the central bar indicates the median, and the whiskers show the 5th to 95th percentile range. nd: not detected. **(B)** Scatter dot plot indicating the relative frequency of *ChAT*^+^ cells within CD4^+^ T cells in patients in circulatory failure “ECMO” (n=33) and healthy control individuals (n=11). Each circle indicates an individual patient. nd: not detected. **(C)** UMAP projection of single cell transcriptomic analysis showing 2703 single CD4^+^ cells from patients in circulatory failure “ECMO” (n=33). Colors denote T cell subsets (see Supplementary Table 4). **(D)** UMAP projection of *ChAT* expression overlaid on T cell subsets from (C). Red denotes *ChAT*^-^ cells and blue denotes *ChAT*^+^ cells. **(E)** Cox proportional hazard rate model for the survival probability for patients (n=32) with low *ChAT*^*+*^*CD4*^*+*^ T cell frequency (Low ChAT Strata, 25^th^ percentile) and high *ChAT*^*+*^*CD4*^*+*^ T cell frequency (High ChAT Strata, 75^th^ percentile) and all other variables (inotrope score and lactate) at their median.

**Table 1:**
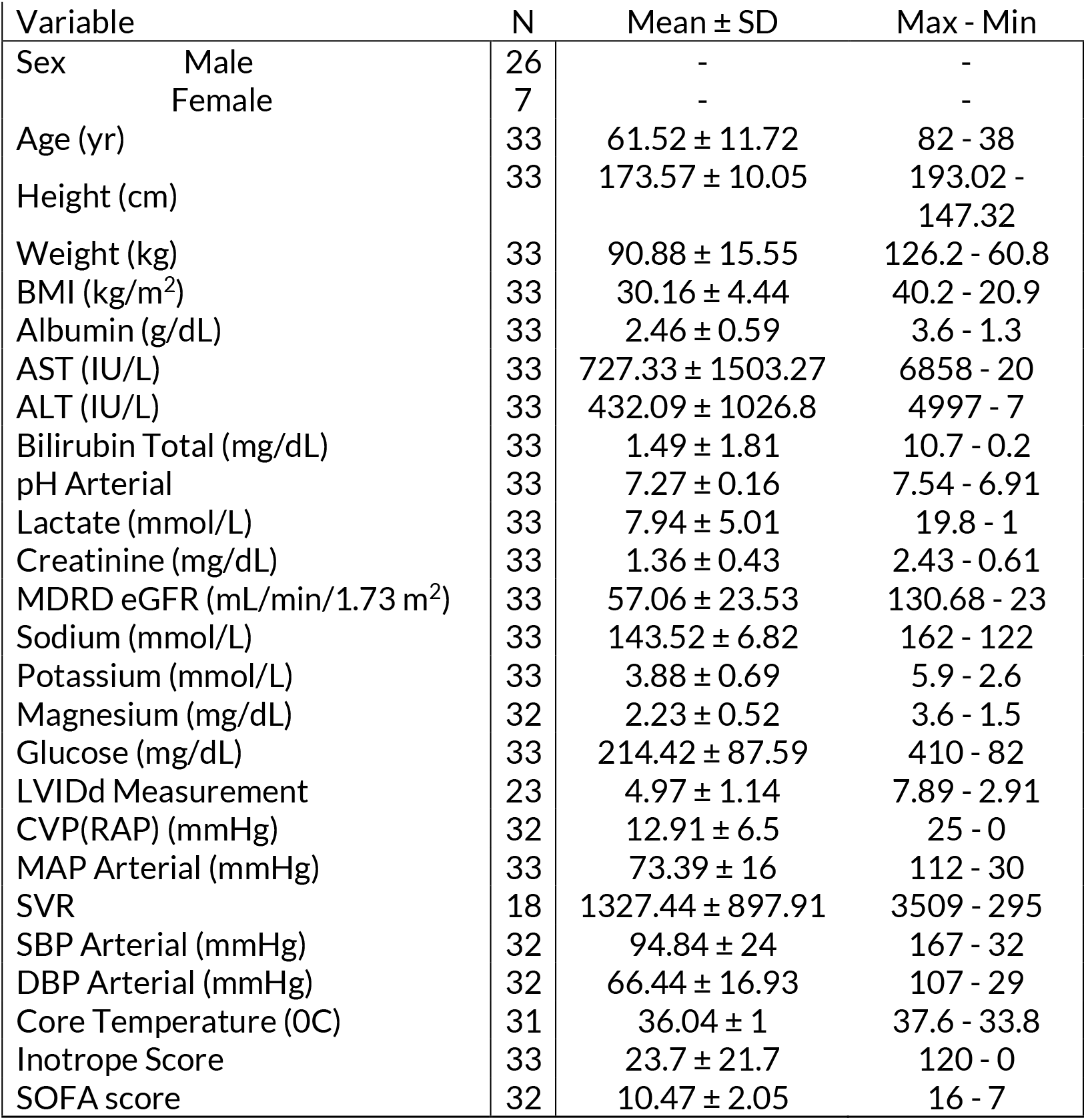
Patient demographics.

In line with our previous findings *in vitro*, we observed that ChAT-expression in human CD4^+^ T cells was not restricted to a specific pre-defined T cell subset in patients with circulatory failure. Unsupervised clustering using Uniform Manifold Approximation and Projection (UMAP) ^30^ of the single cell transcriptomic data showed that all cell clusters (Figure 4C, D) and pre-defined T cell subsets (Figure S5A, Table S4) contained *ChAT*^+^ cells. There was no difference in the proportion of *ChAT*^+^ cells between the different CD4^+^ subtypes (Holm adjusted p-values for pairwise comparisons of proportions: 0.3-1.0), indicating that multiple *CD4*^+^ T cell subsets in blood within this cohort express *ChAT*.

### The relative frequency of CD4^+^ ChAT^+^ T cells in blood correlated with survival

Considering the reported correlation of blood acetylcholine levels and outcome in the critically ill, we next studied ChAT^+^ CD4^+^ T cells in context of the clinical data. There was a weak negative correlation (Spearman r (r_S_) = -0.35, p=0.05) between relative *ChAT*^*+*^*CD4*^*+*^ T cell frequency and maximum blood lactate (Figure S5B), a clinical measurement associated with insufficient organ perfusion and increased mortality in the critically ill ^31,32^. Thus, we reasoned that a low proportion ChAT^+^CD4^+^ T cells in blood might be a risk factor for adverse outcomes. Accordingly, we investigated whether *ChAT*^*+*^*CD4*^*+*^ T cell relative frequency in blood correlated to survival in this cohort. To study this, we utilized a Cox proportional hazard rate model with several covariates. Our optimized model included ChAT^+^ cell relative frequency, sex, inotrope score and lactate as significant explanatory variables. For each percentage point increase in ChAT^+^ cell relative frequency, the hazard ratio in our cohort decreased by a factor 0.97, all other variables equal (95% CI 0.94-0.99, p-value 0.01) (Table S5). Hence, patients in this cohort with a low proportion of ChAT^+^ cells (25-percentile) had almost a six-fold (5.9) higher hazard rate for 30-day mortality compared to patients with high proportion of ChAT^+^ cells (75-percentile), all other variables equal. The corresponding survival probability for *ChAT*^*+*^ cell relative frequency (at the 25/75-percentile) is represented in Figure 4E, with all other variables (inotrope score and lactate) at their median. Together, these observations show a correlation between mortality and the *ChAT*^*+*^*CD4*^*+*^ T cell relative frequency in blood and suggest that low *ChAT*^*+*^*CD4*^*+*^ T cell relative frequency is a risk factor for death in this cohort.

While the correlation survival observed here should be interpreted with some caution considering the small sample size and the diverse clinical background of the included patients, the effect size was large and the observation is interesting. It is conceivable that local vasodilatation and entry of antigen-specific T cells mediated by ChAT^+^ T cells, as observed in mice ^3^ and supported by the findings here, may improve organ perfusion and pathogen clearance, and contribute to recovery in the critically ill. It will be important to further study the mechanisms and effects of ChAT^+^CD4^+^ T cell-mediated vasodilation and their pathophysiological role in human disease ^12^.

In conclusion, the identification and characterization of ChAT^+^CD4^+^ T cells here provides evidence that human T cell ChAT is regulated by activation and GATA3. ChAT^+^CD4^+^ T cells released ACh that promoted arterial relaxation *in vitro*, and their relative frequency correlated with survival in a cohort of critically ill patients. Considering the important physiological role of ChAT^+^ T cells observed in experimental animals, the identification of human ChAT^+^CD4^+^ T cells lays the groundwork for the study of another hitherto unexplored endogenous mechanism in immune cell-tissue interactions. An improved understanding of this physiology may provide opportunities to design novel therapies that regulate local infiltration of immune cells at target sites.

## Materials and methods

### Isolation of human peripheral blood mononuclear cells and T cells

Whole blood from healthy individuals were purchased from the Immunology and Transfusion Medicine department at Karolinska University Laboratory. Peripheral blood mononuclear cells (PBMC) were isolated, cryopreserved at -80°C, and transferred to liquid nitrogen for extended storage. T cells were isolated using the human Pan T cell Isolation Kit (Miltenyi Biotec) and naive CD4^+^ T cell using the Naive CD4^+^ T Cell Isolation (Miltenyi Biotec). T cells were activated using ImmunoCult™ Human CD3/CD28 T Cell Activator antibodies (StemCell Technologies).

### T cell activation and differentiation

Cryopreserved healthy human CD3^+^ T cells or CD4^+^ T cells were activated and maintained in culture for 0-14 days. CD4^+^ T cell differentiation was carried out using ImmunoCult™ Human Th1 Differentiation Supplement (StemCell Techno.), ImmunoCult™ Human Th2 Differentiation Supplement (StemCell Techno.), and ImmunoCult™ Human Treg Differentiation Supplement (StemCell TechnoRNA was isolated at set time points after activation using the RNeasy Mini (Qiagen). cDNA was generated using the High-Capacity RNA-to-cDNA Kit (ThermoFisher), and analyzed by qPCR for mRNA levels using TaqMan Universal PCR Master Mix (ThermoFisher).

### Flow cytometry analysis

Activated T cells were stained using antibodies against CD25 (ThermoFisher,) and with LIVE/DEAD™ Fixable Aqua Dead Cell Stain (ThermoFisher), and fixed using the Foxp3/Transcription Factor Staining Buffer Set (eBioscience). Stained cells were analyzed using the BD FACSVerse system (BD Biosciences) and FlowJo software v.10 (FlowJo).

### ACh release

Activated (96 h) healthy human T cells were stained for viability using DAPI (ThermoFisher). Cells were isolated based on size using the Influx fluorescence-activated cell sorter (BD Biosciences), sorted into PBS in the presence of pyridostigmine bromide (PBr) and incubated for 30 minutes. Alternatively, activated (0-120 h) healthy human T cells were incubated in 200 µL PBS+PBr for 20 minutes. The cell-free conditioned PBS supernatant was snap frozen and stored at -80°C until mass spectrometry analysis.

### Liquid chromatography tandem mass spectrometry

All chromatography measurements were blinded. Concentrations of ACh in the supernatants were measured by ultra-high performance liquid chromatography tandem mass spectrometry (UHPLC-MS/MS). The calibration standards were prepared in PBS, the calibration curve was linear in the range of 0.1 – 256 nM (R2 = 0.999), the estimated limit of detection (LOD) was 0.05 nM, the lower limit of quantification (LLOQ) was 0.15 nM.

### Myograph

All myograph measurements were performed blinded. Activated primary human T cells were incubated in PBS in the presence of PBr for 15 minutes to make conditioned PBS. Descending thoracic aortas from C57BL6 mice (Janvier) were dissected in ice-cold physiological salt solution and arterial rings (2 mm) were mounted onto the pins of multi wire myograph system (Model 620M; Danish Myo Technology, Denmark). The chambers were filled with PSS solution (37 °C, pH 7.4) aerated with carbogen (95% O2; 5% CO2). Isometric tension was recorded with Powerlab system (Powerlab 4/30). After a 45 min equilibration a loading force of 6 mN was added to the vessel. After another 45 min equilibrating period, vessels were pre-constricted with phenylephrine. Endothelial function was accessed by acetylcholine (ACh) dose-response curve measurements. The vessels were washed and equilibrated, pre-constricted with phenylephrine, and exposed to vehicle (PBS+PBr) or pools of cell-free T cell-conditioned PBS, with and without pre-treatment of the vessel with atropine.

### Measurement of NO species using DAF-FM fluorescence

All DAF measurements were blinded. Activated (96 h) primary human T cells were incubated in PBS in the presence of PBr for 15 minutes to make conditioned PBS. Human Umbilical Venous Endothelial Cells (HUVEC; pooled from 3 individual donors; LONZA) were starved for 6 h, washed in PBS, and exposed to vehicle (PBS) or the cell-free conditioned PBS at 10% or 2%, in the presence of 4-amino-5 methylamino-2’
s,7’-difluorofluorescein (DAF-FM) diacetate (Sigma, D1946). DAF fluorescence intensities were measured every 5 minutes for a total of 60 minutes using a SpectraMax iD3 (Molecular Devices). All experiments were repeated at least four times per batch.

### Nitric Oxide Synthase (NOS) activity

All NOS activity measurements were blinded. Activated primary human T cells or primary human CD4^+^ T cells were incubated in PBS in the presence of PBr for 15 minutes to make conditioned PBS. Human EA.hy926 endothelial cells (ATCC CRL-2922) were washed with PBS and exposed to vehicle (PBS+PBr) or the cell-free conditioned PBS, with and without Atropine for 10 minutes. Protein concentration in endothelial cell lysates were measured using bicinchoninic acid assay (BCA) Protein Assay Kit (Abcam, Cambridge, UK). NOS production was measured indirectly by Griess reagent in samples using Nitric Oxide Synthase (NOS) Activity Assay (Abcam, Cambridge, UK) with the addition of Protease Inhibitor Cocktail (Abcam). The assays were performed per manufacturer’s instructions. The absorbance was measured using VersaMax microplate reader (Molecular Devices, California, US).

### FACS cell isolation, cDNA pre-amplification and quantification

Activated T cells were stained using antibodies against CD4 (BD Pharmingen), CD8 (BD Pharmingen) or CD25 (BD). For each population of interest, 50 cells for bulk analysis or one cell for single cell analysis, were isolated using the Influx fluorescence-activated cell sorter (BD Biosciences), sorted into a 96-well plate containing 5 µL Fluidigm master mix per well. The hydrolysis probes used for cDNA amplification during the Fluidigm and qPCR experiments are found in Table S1. Stained cells were analyzed using the FlowJo software v.10 (FlowJo). cDNA was pre-amplified using the Veriti™ 96-Well Thermal Cycler (Applied Biosystems) or S1000™ Thermal Cycler (Bio-Rad). Single cell qPCR was performed blinded according to manufacturer protocol using the high-throughput BioMark Real-Time PCR system (Fluidigm). Bulk mRNA was analyzed by qPCR using TaqMan Universal PCR Master Mix (ThermoFisher).

### Transcription factor binding site identification

ChAT 5’-regulatory region (chr10:49612453-49615546) was scanned for the presence of potential transcription factor binding sites using PWMTools^26^. JASPAR-derived binding matrixes^33^ were used for scanning with the cutoff of p<0.05. The potential binding sites discovered by PWMTools were filtered to remove site with more than one mutation. Locations of TF binding sites on ChAT 5’-regulatory region were visualized using R/Bioconductor package Gviz^34^.

### Lentiviral transduction

Human naive CD4^+^ T cells were activated, maintained in culture for 10 days. Lentiviral transfection was carried out using spinoculation with either scrambled shRNA (Santa Cruz) or shRNA targeting GATA3 mRNA (Santa Cruz). Puromycin (Santa Cruz) was used to select for transfected cells.

### Patient sample collection and analysis

The VA-ECLS study was approved by the Spectrum Health Review Board. All patients or their legally authorized representative gave informed consent to participate. All samples and data were anonymized before analysis. Whole blood was collected in EDTA Vacutainers. PBMCs and plasma was isolated and cryopreserved at -80°C until analysis.

Plasma from each patient sample was analyzed using the Bio-Plex Pro™ Human Cytokine 17-plex Assay kit (Bio-Rad). Cytokine expression was measured using the Luminex xMAP® (Luminex Corporation) fluorescent bead-based technology in a capture/detection sandwich immunoassay format and read on a Bio-Plex 200 system using Bio-Plex Manager software (v.6.1). Cytokine concentrations were calculated using an eight-point standard curve.

### Single cell encapsulation, reverse transcription and sequencing

Cryopreserved PBMCs from each patient sample were washed and stained for viability using SYTOX Green (Thermo Fisher Scientific). Samples were sorted on a BD Influx flow cytometer into Phosphate Buffered Saline (PBS). Cells were encapsulated with barcoded beads and processed using the inDrop platform (1CellBio). The resulting cDNA was cleaned using MinElute columns (Qiagen). Second strand synthesis was performed using the NEBNext Ultra II second strand synthesis kit (NEB). *In vitro* transcription was performed using the NEB HiScribe High Yield RNA synthesis kit. The entire process was performed according to protocols supplied by the manufacturer. Library integrity and fragment size were confirmed by microfluidic gel electrophoresis Bioanalyzer (Agilent) prior to sequencing. Libraries were sequenced on a NovaSeq 6000 sequencer (Illumina) using the S2 100 cycle kit.

### Data processing

The sequencing data was aligned to the human genome (assembly GRCh38) and unique feature counts obtained using the software pipeline developed by the inDrop manufacturer (https://github.com/indrops/indrops). Technical dropouts were imputed using the ALRA algorithm ^35^. The raw count data was then filtered and normalized for library size using version 3 of the Monoclec toolset ^36^ for the R statistical analysis platform. Expression data for healthy donors was obtained from 10x Genomics publicly available Single Cell Gene Expression and Single Cell Immune Profiling datasets (Table S6). UMAP batch correction was performed according to Korsunsky et.al ^37^.

### Study approval

The VA-ECLS study was approved by the Spectrum Health Review Board. All patients or their legally authorized representative gave informed consent to participate. All samples and data were anonymized before analysis. Animal experiments were performed in accordance with guidelines from the Swedish National Board of Laboratory Animals under protocols approved by the Animal Ethics Review Board of Stockholm, Sweden (No. N139/15).

## Author Contributions

LT and PSO conceived, designed and planned the study. SJ and EJK designed and performed the VA-ECLS clinical study. LT, ALG, VS, ZZ, MW, ASC, SS, FHW performed experiments. LT, ALG, ZZ, ASC, DME, ME, MC, PSO analyzed experimental data. LT, EJK, MW, HH, MG, SJ, and PSO analyzed the data from the VA-ECLS clinical study. LT, ALG, EJK, VS, ASC, ME, JK, SGM, MC, SJ, PSO interpreted the results. The scRNAseq was performed in the group of SJ. EJK analyzed the single cell RNA sequencing data. LT, ALG, PSO wrote the manuscript. LT, AF, EJK, EW, ME, JK, SGM, MC, SJ, PSO critically reviewed or revised the manuscript. All authors contributed to the material and methods, reviewed the drafts and final version of the manuscript, and agreed with its content and submission.

## Acknowledgements

We would like to thank the patients and their families for participating in this study, Jennifer Schuitema for overseeing recruitment, consent, and collection of blood samples, the ECMO nursing staff for collecting blood samples and processing plasma after-hours, and David Chesla and Donald Daley from the Spectrum Health Biorepository for sample processing. The authors thank the Van Andel Genomics Core for providing sequencing facilities and services, Joseph Faski for performing the Fluidigm Biomark run, and Emily Eugster for the creation of the cDNA libraries as part of the scRNAseq. This study was supported by grants from the MedTechLabs (2019), ALF Project Funds (20170199, 20180502), Knut and Alice Wallenberg foundation (2014), The Swedish Research Council (2017-03366, 2020-04443), the Swedish Heart-Lung Foundation (20200827, 20190672) to PSO, the Lars Hiertas Minne Foundation (FO2018-0494), the Gösta Fraenckel Foundation (2019-01096), the Loo and Hans Osterman Foundation (2019-01408), and Foundation for Geriatric Diseases (2019-01337) to LT, and the Richard and Helen DeVos Foundation (2014-JOVINGE) to SJ. Conflict of interest statement: PSO is a shareholder of Emune AB. The authors have no additional financial interests.

## SUPPLEMENTAL MATERIAL

### Supplemental Figures

**Figure S1:**
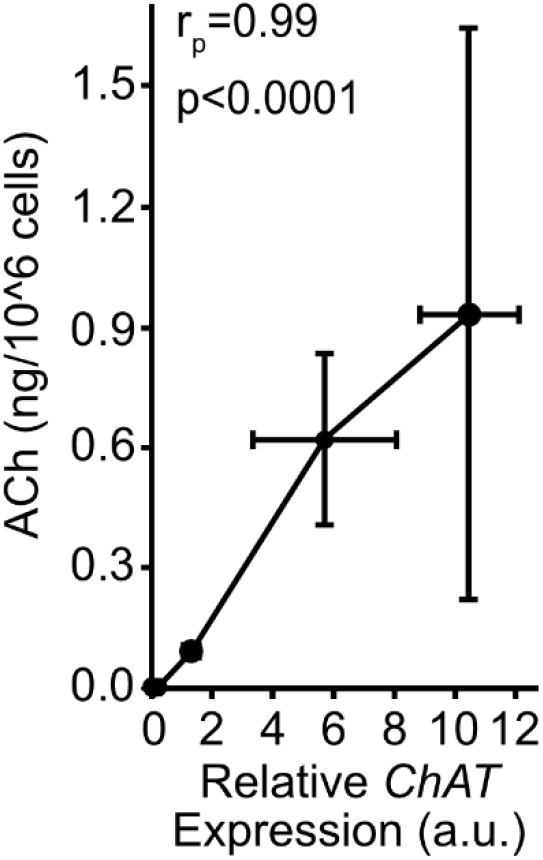
Correlation between *ChAT* expression and ACh release from human T cells following activation. Primary human T cells were activated using antibodies against CD3 and CD28 and harvested at 0, 24, 48, 72, 96, and 120 h. *ChAT* mRNA was quantified as in Figure 1A and supernatants analyzed for ACh using mass spectrometry as per Figure 1D. qPCR data are normalized to reference genes *PPIA* and *B2M*, all time points are relative to *ChAT* expression at 72 h, and represented as mean ± SEM. Mass spectrometry values are represented as mean ACh in ng ± SEM per 10^6^ T cells. (Pearson r (r_p_) r=0.99, p<0.0001).

**Figure S2:**
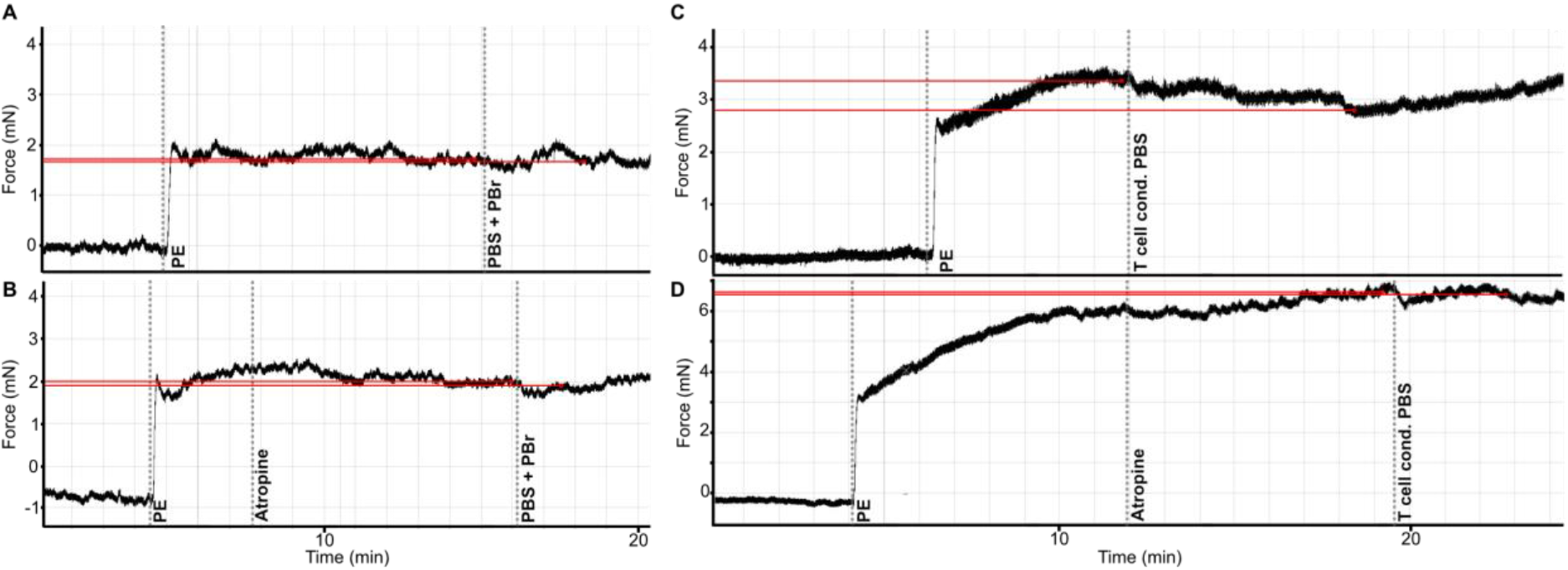
Representative tracings of the vascular relaxation mouse aortas pre-contracted with phenylephrine (PE) and exposed to A-B, vehicle, with or without pre-treatment with atropine (1.7 µM) and C-D, T cell conditioned PBS, with or without pre-treatment with atropine (1.7 µM).

**Figure S3:**
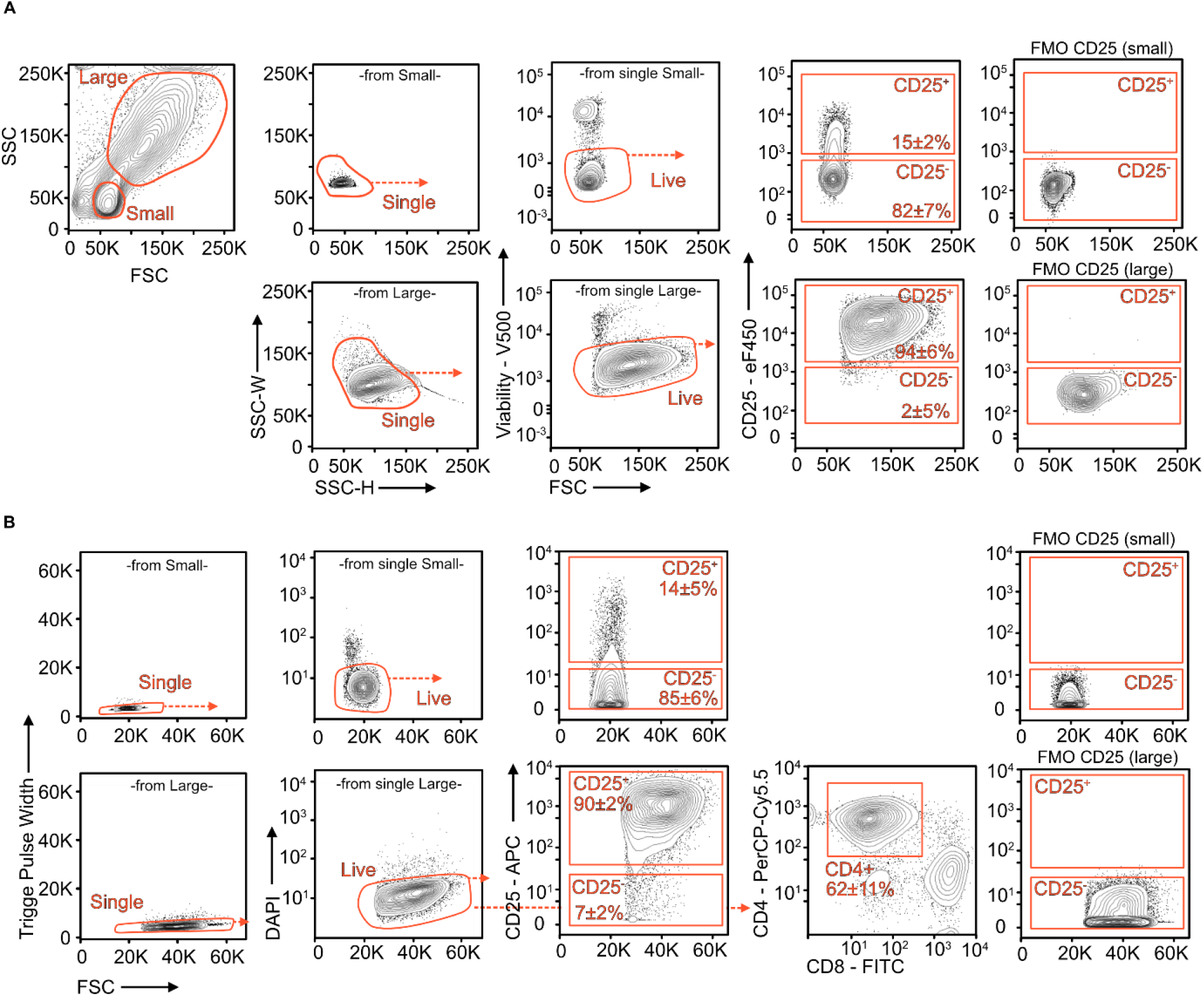
Representative plots of flow cytometry gating and analysis of primary human T cells activated for (a) 72 h (n=8) or (b) 96 h (n=6) (Primary gate: Fig. 1B). Cells were selected based on size: Small (SSC/FSC Low) or Large (SSC/FSC High), single cells, viability/DAPI, and interrogated for CD25 and CD4 expression. Positive and negative gates were set based on fluorescence minus one (FMO) controls (n=4). Gates are shown as median percentage ± SEM.

**Figure S4:**
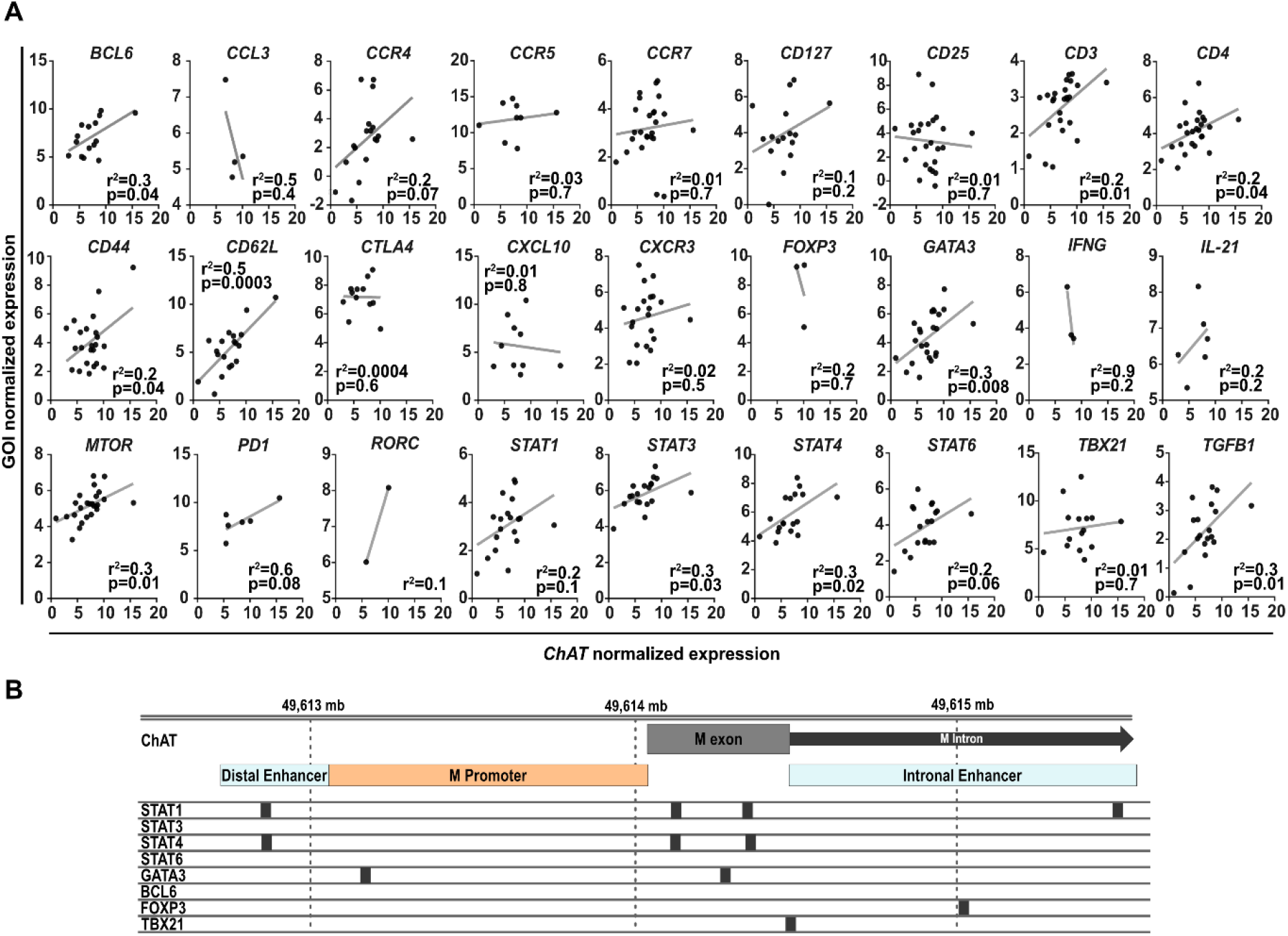
**A)** Linear regression analysis of *ChAT* expression and genes enriched in distinct immune cell phenotypes within single *CD4*^+^ *ChAT*^+^ T cells (n=25). Activated *CD4*^+^ single T cells were isolated by FACS and mRNA levels measured using the Fluidigm Biomark single cell analysis. Data points represent threshold values normalized to the reference gene *PPIA*. Measurements below detection were excluded from the graph. Correlations were calculated using linear regression. *p<0.05, **p<0.01, ***p<0.001. **B)** Schematic representation map of potential transcription binding motifs within the ChAT promoter region for *STAT1, STAT4, TBX21, FOXP3, GATA3, STAT3, STAT6, and BCL6* as determined using PWMTools. JASPAR-derived binding matrixes were used for scanning with the cutoff of p<0.05. Locations of TF binding sites on ChAT 5’-regulatory region were visualized using R/Bioconductor package Gviz. Black boxes indicate approximate predicted binding motif.

**Figure S5:**
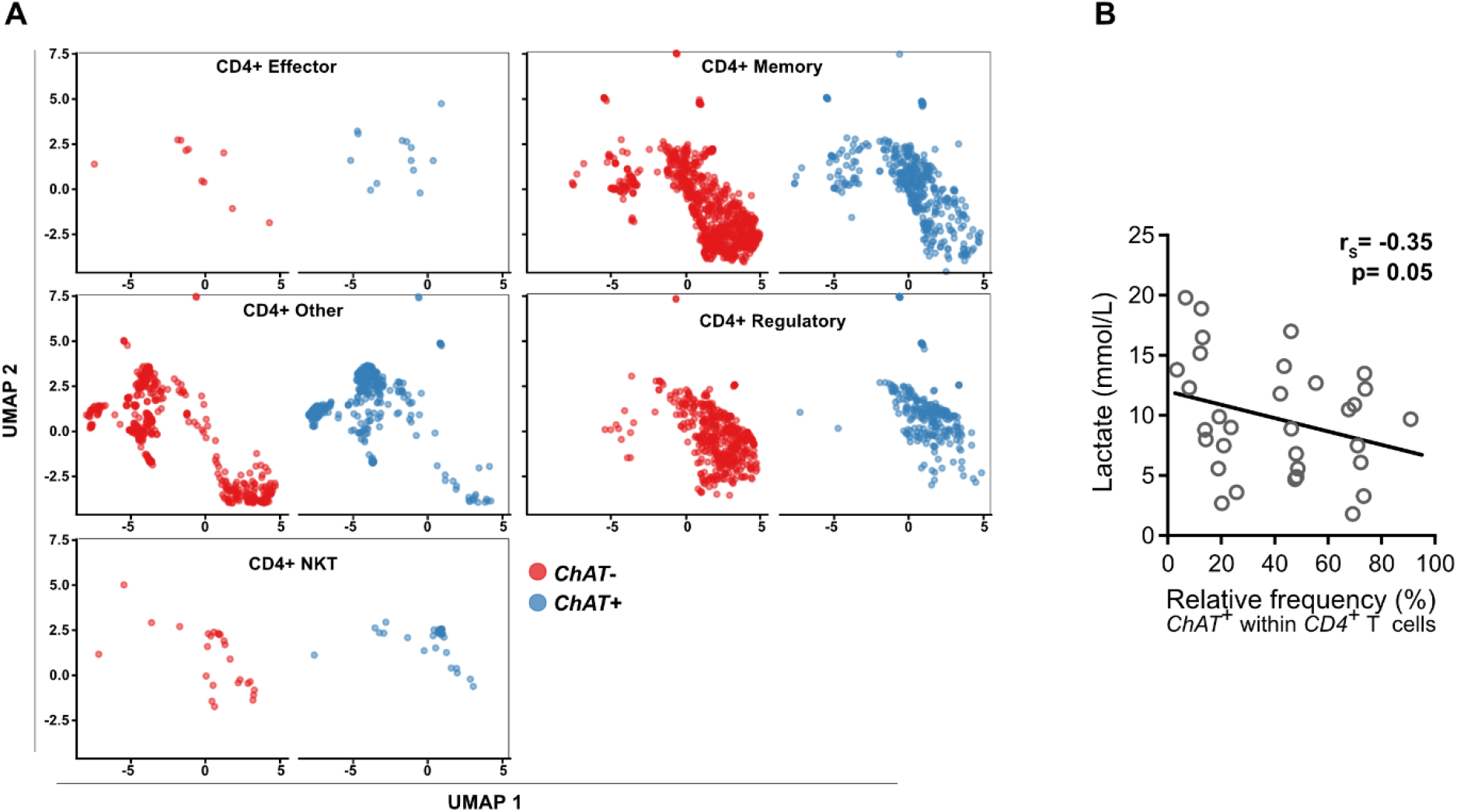
**A)** UMAP projection of single cell transcriptomic analysis of 2703 single *CD4*^+^ cells from critically ill patients (n=33) separated by T cell subset maker expression (Supplementary Table 5). *ChAT*^-^ and *ChAT*^+^ cells are designated by red and blue color, respectively. **B)** Correlation between lactate levels and the relative frequency of *ChAT*^+^ cells of verified *CD4*^+^ T cells in critically ill patients with circulatory shock (n=32). Each circle shows individual patients. Spearman r (r_S_) = -0.35, p=0.05.

### Supplemental Tables

**Table S1:**
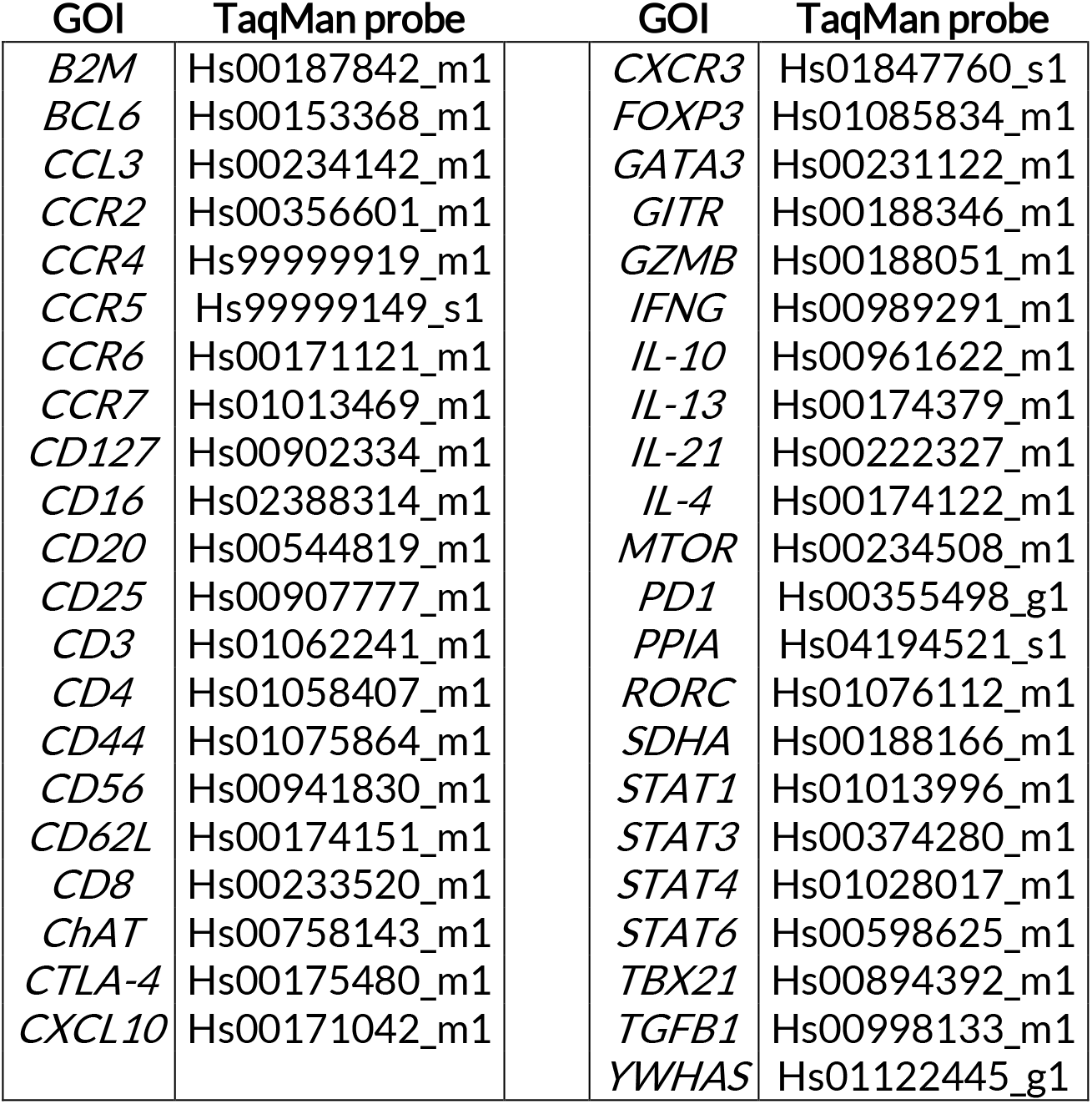
TaqMan hydrolysis probes used in Fluidigm gene expression analysis and qPCR analysis ^25^.

**Table S2:** Cause of admission of patients in imminent need of Veno-Arterial Extra Corporeal Life Support (VA-ECLS).

| Cause of admission | N |
| --- | --- |
| Coronary Artery Disease | 9 |
| Congestive Heart Failure | 4 |
| Arrhythmia | 4 |
| Thrombosis | 2 |
| Dissection or aneurysm | 2 |
| Cardiomyopathy | 1 |
| Aortic Valve Stenosis | 1 |
| Pulmonary Fibrosis | 4 |
| Respiratory Failure | 1 |
| Sepsis | 2 |
| Postoperative Shock | 1 |
| Trauma | 1 |
| Seizure | 1 |

**Table S3:** Plasma cytokine levels in the study population.

| Cytokine | Detectable in patients: | Mean, SD, and minimum-maximum, serum cytokines levels (pg/mL) | Mean, SD, and minimum-maximum, serum cytokines levels of young healthy volunteers (Age < 45) (pg/mL) <sup>29</sup> |
| --- | --- | --- | --- |
| IL1β | 31/31 | 3.0 ± 3.7 (0.43–16) | 2.04 ± 4.9 (0.17–24) |
| IL2 | 7/31 | 1.1 ± 4.5 (0.0–24.9) | 5.1 ± 2.3 (2.8–18) |
| IL4 | 28/31 | 1.6 ± 2.4 (0.0–9.7) | - |
| IL5 | 17/31 | 1.4 ± 2.5 (0.0–12) | - |
| IL6 | 31/31 | 2266 ± 7150 (1.6–38537) | 2.9 ± 6.5 (0.16–38) |
| IL7 | 29/31 | 3.9 ± 3.6 (0.0–16) | - |
| IL8 | 31/31 | 68.4 ± 102 (2.9–387) | 24 ± 30 (4.2–133) |
| IL10 | 30/31 | 138 ± 226 (0.0–1039) | 1.3 ± 3.1 (0.01–20) |
| IL12 | 30/31 | 6.8 ± 5.2 (0.0–22) | - |
| IL13 | 20/31 | 2.3 ± 4.3 (0.0–17) | - |
| IL17 | 30/31 | 48 ± 43 (0.0–142) | 6.5 ± 7.4 (1.6–38) |
| G-CSF | 31/31 | 654 ± 2152 (8.6–11416) | 15 ± 13 (0.03–76) |
| GM-CSF | 23/31 | 48 ± 67 (0.0–252.8) | 41 ± 109 (0.5–728) |
| IFNγ | 22/31 | 47 ± 77 (0.0–249.9) | 13 ± 23 (0.14–127) |
| MCP | 29/31 | 217 ± 277 (0.0–1074) | 214 ± 101 (28–668) |
| MIP-1b | 31/31 | 98 ± 85 (19.32–348.5) | 41 ± 39 (3.2–227) |
| TNFα | 31/31 | 39 ± 69 (0.787–277.6) | 3.2 ± 4.04 (0.93–27) |

**Table S4:** Classification of PBMC subsets based on scRNASeq expression in the UMAP analysis for naive, memory, effector and regulatory CD4^+^ T cells.

|  | CD3 | CD4 | CD8 | CD2 | CD57 | CD25 | FOXP3 | CD56 |
| --- | --- | --- | --- | --- | --- | --- | --- | --- |
| CD4 Other | + | + | - | - |  | +/-<sup>a</sup> | +/-<sup>a</sup> | - |
| CD4 Memory | + | + | - | + | - | +/-<sup>a</sup> | +/-<sup>a</sup> | - |
| CD4 Effector | + | + | - | + | + | +/-<sup>a</sup> | +/-<sup>a</sup> | - |
| CD4 Regulatory | + | + | - |  |  | + | + | - |
| CD4 NKT | + | + | - |  |  |  |  | + |

<sup>a</sup>Not CD25/FOXP3 double positive

**Table S5:** Hazard ratio and 95% confidence interval of mortality in the study population adjusted for co-variates (n=32).

| Covariate | Hazard ratio | 95% Confidence interval | P-value |
| --- | --- | --- | --- |
| *ChAT*<sup>+</sup> frequency | 0.965 | (0.939, 0.992) | 0.012 |
| Inotrope score | 1.043 | (1.007, 1.080) | 0.017 |
| Sex | 0.258 | (0.069, 0.964) | 0.044 |
| Lactate | 1.231 | (1.060, 1.429) | 0.006 |

**Table S6:** Publicly available Single Cell Gene Expression and Single Cell Immune Profiling datasets of healthy individuals.

| Data set name: | Number of Cells (after QC): | Number of CD4<sup>+</sup> T cells | Single Cell Gene Expression Dataset source: |
| --- | --- | --- | --- |
| fresh_68k_pbmc_donor_a | 69941 | 2307 | Cell Ranger 1.1.0 |
| frozen_pbmc_donor_b | 7976 | 210 | Cell Ranger 1.1.0 |
| frozen_pbmc_donor_c | 9773 | 395 | Cell Ranger 1.1.0 |
| vdj_pbmc_5gex | 9272 | 1027 | Cell Ranger 2.2.0 |
| vdj_pbmc_v2_10k | 10691 | 1899 | Cell Ranger 4.0.0 |
| pbmc_10k_v3 | 11934 | 1626 | Cell Ranger 3.0.0 |
| pbmc33k | 33700 | 1505 | Cell Ranger 1.1.0 |
| 5k_pbmc_v3 | 5053 | 981 | Cell Ranger 3.0.2 |
| SC3_v3_NextGem_SI_PBMC_10K | 12848 | 1857 | Cell Ranger 4.0.0 |
| vdj_v1_hs_pbmc3_5gex_protein | 7441 | 1145 | Cell Ranger 3.1.0 |
| Parent_SC5_PBMC | 6795 | 1228 | Cell Ranger 4.0.0 |

